# Discovery and Validation of Context-Dependent Synthetic Mammalian Promoters

**DOI:** 10.1101/2023.05.11.539703

**Authors:** Adam M. Zahm, William S. Owens, Samuel R. Himes, Kathleen E. Rondem, Braden S. Fallon, Alexa N. Gormick, Joshua S. Bloom, Sriram Kosuri, Henry Chan, Justin G. English

## Abstract

Cellular transcription enables cells to adapt to various stimuli and maintain homeostasis. Transcription factors bind to transcription response elements (TREs) in gene promoters, initiating transcription. Synthetic promoters, derived from natural TREs, can be engineered to control exogenous gene expression using endogenous transcription machinery. This technology has found extensive use in biological research for applications including reporter gene assays, biomarker development, and programming synthetic circuits in living cells. However, a reliable and precise method for selecting minimally-sized synthetic promoters with desired background, amplitude, and stimulation response profiles has been elusive. In this study, we introduce a massively parallel reporter assay library containing 6184 synthetic promoters, each less than 250 bp in length. This comprehensive library allows for rapid identification of promoters with optimal transcriptional output parameters across multiple cell lines and stimuli. We showcase this library’s utility to identify promoters activated in unique cell types, and in response to metabolites, mitogens, cellular toxins, and agonism of both aminergic and non-aminergic GPCRs. We further show these promoters can be used in luciferase reporter assays, eliciting 50-100 fold dynamic ranges in response to stimuli. Our platform is effective, easily implemented, and provides a solution for selecting short-length promoters with precise performance for a multitude of applications.

## Introduction

Transcription factors (TFs) bind to specific DNA sequences known as transcription response elements (TREs) to initiate gene transcription^1–4^. Identical TREs can be found at multiple loci across the genome, serving as platforms to facilitate coordinated transcription programs for gene network activation^5,6^. The binding of TFs to TREs and their interaction with enhancers and other distal regulatory elements define how individual genes are regulated transcriptionally^7^.

The TF-TRE mechanism has long been harnessed to create engineered systems, where active TFs in the cell drive transcription of user-selected genetic material by binding to TREs in genome-derived or synthetic promoters. Promoters responding to specific cellular signaling events by modulating transgene expression in a reproducible manner are crucial for well-established applications (e.g. reporter assays) and emerging technologies, such as pluripotent stem cell lineage-control networks and regulatory control of human CAR-T cells *in vivo*^8,9^. Existing synthetic promoters often consist of TREs arrayed immediately upstream of an inactive or weakly functional minimal promoter. Transcription factor binding to the response elements activates the minimal promoter, initiating transcription. Due to their short length relative to endogenous promoter regions, synthetic promoters are advantageous for many applications, including those utilizing vectors with limited cargo sizes. However, off-the-shelf availability is extremely limited, often necessitating the in-house development of new promoters with desired characteristics (e.g. high dynamic range, defined basal activity, short length, etc.). The ability to quickly screen genome-derived or synthetic promoters suitable for specific applications at large scale will accelerate molecular tool development and deployment.

Technologies such as self-transcribing active regulatory region sequencing (STARR-seq) and massively parallel reporter assays (MPRAs) have been used to interrogate the effects of DNA sequence on gene regulation at extremely high throughput^10–16^. Here, we present a MPRA library for the rapid identification of synthetic promoters that serve as downstream transcriptional readouts of specific upstream signaling events in mammalian cells. 6184 synthetic promoters of less than 250 bp in length, containing candidate TREs of 229 human and mouse TFs, are represented by over 3.6 million barcoded plasmids. We demonstrate that single replicate transient transfections of this library can provide reliable transcription rate estimates for episomal synthetic promoters across a range of stimuli, from heavy metal toxicity to G protein coupled receptor (GPCR) activation.

## Results

### Identification of functional synthetic promoters using a massively parallel reporter assay

Coupling promoter activity to the production of barcoded mRNAs affords the ability to quantify activity in response to stimuli in a sensitive and high-throughput manner via next-generation sequencing (NGS)^11^. We developed a barcoded plasmid library (‘TRE-MPRA’) of synthetic promoters composed of TF binding motifs derived from the DNA position weight matrices (PWMs) for hundreds of human and mouse TFs identified via HT-SELEX^17^. We reasoned that TREs based on DNA sequences bound by TFs *ex vivo* would produce superior synthetic promoters compared to sequences based on genomic footprints and removed from their native chromatin context. Four copies of a given binding motif were arranged in six configurations (TRE units) and positioned immediately 5’ to one of three minimal promoters to create short, synthetic promoters (hereafter ‘promoters’) driving expression of a protein coding transcript (Luc2) with a barcoded 3’ UTR (Fig. 1, S1). We also included hundreds of negative control promoters containing TREs of scrambled sequences in the library. Three independent plasmid library preparations from distinct liquid cultures were sequenced on separate flow cells and showed nearly identical barcode representation per promoter, as well as highly correlated barcode reads per million (Fig. S2). The mean and median barcode representation for promoters in a representative plasmid library preparation were 82 and 65, respectively. Of the 6318 promoters we designed, a total of 6184 (98%) were detected in our plasmid preparations.

**Figure 1.**
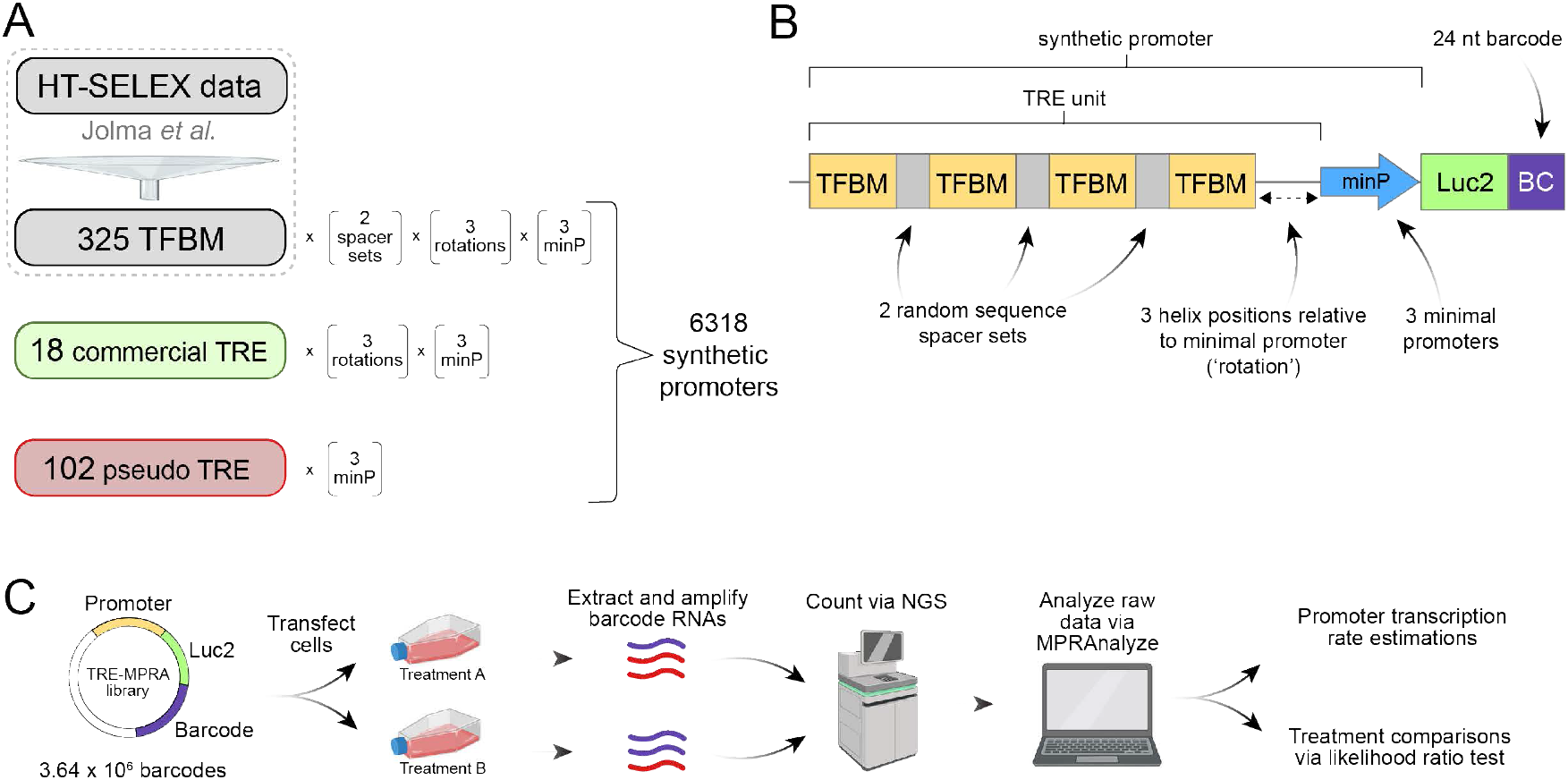
TRE-MPRA overview. **A)** Transcription factor binding motifs (TFBMs) included in the TRE-MPRA library were derived from the HT-SELEX dataset of Jolma *et al*.^17^ Candidate synthetic promoters were designed by combining homotypic TFBMs and one of three minimal promoters in multiple configurations. Commercially available TREs, as well as negative control promoters (‘pseudo’) were also included in the TRE-MPRA library. **B)** Four copies of a given TFBM were oriented on alternating sides of the DNA double helix using sets of random nucleotides and rotated along the helix relative to the minimal promoter (TRE unit). TRE units combined with minimal promoters (‘promoters’) regulate the expression of a Luc2 CDS containing 24 nucleotide barcodes in the 3’ UTR. Barcodes were mapped to associated TRE units using next-generation sequencing (NGS) during library preparation. **C)** Barcoded RNAs were extracted from transfected cells and sequenced via NGS alongside the input TRE-MPRA plasmid libraries. Resulting barcode counts were analyzed via MPRAnalyze^18^ to produce estimates of promoter transcription rates and to compare activities between treatments.

We first benchmarked this library by transfecting the human HEK293 cell line and culturing the cells in serum-free media to establish baseline transcriptional activities of each promoter. We quantified barcoded mRNA levels via NGS as a readout of promoter transcriptional activity and then estimated transcription rates using barcode frequencies in the input plasmid pool and RNA fractions with MPRAnalyze^18^. Each of two independent experiments, both comprising four independent replicates of transfected cells and sequenced on separate flow cells, showed a range of transcriptional rate estimates greater than 200-fold across all promoters (Fig. S2). Furthermore, rate estimates were highly correlated across the independent experiments (Spearman’s ρ = 0.952, p < .0001) (Fig. S2). Across the population of TRE units, we observed a pronounced effect of the paired minimal promoter on baseline transcription rates, whereas the spacer sequences between TF binding motifs and the distance between the TRE unit and minimal promoter did not globally alter transcription (Fig. S3).

Next, we transfected HEK293 cells with the TRE-MPRA library and subsequently treated the cells with fetal bovine serum (FBS) or forskolin in triplicate. Cellular responses to serum and forskolin are classically associated with the serum response element (SRE) and cAMP response element (CRE), respectively^19–21^. Differential promoter activities relative to untreated cells transfected with the library were determined using MPRAnalyze. As expected, serum treatment elevated expression from SRE promoters, while forskolin activated CRE promoters (Fig. 2A). Transcriptional activities from promoters containing one of three human thyroid hormone receptor beta (THRB) motifs (THRB-1) were significantly elevated in cells treated with FBS, whereas motifs of highly-similar sequence did not alter transcription in response to FBS (Fig. 2A, B). We also observed increased activities from promoters containing the murine v-maf musculoaponeurotic fibrosarcoma oncogene family, protein B (Mafb) binding motif following stimulation with forskolin (Fig. 2A). Of note, the THRB-1 motif is highly similar in sequence to the CArG box of SRE bound by serum response factor (SRF) and thus may provide a readout of SRF activity rather than THRB (Fig. 2B)^21^. Likewise, the Mafb motif (TGCTGACGTAAGCA) in the TRE-MPRA library contains a sequence very similar to the cAMP responsive element (TGACGTCA) and so may be activated by cAMP responsive element binding protein 1 (CREB1) as opposed to MAFB^20^. Importantly, the response of these promoters was independent of barcode abundance in the plasmid library (Fig. S4A). Both treatment conditions showed differential effects of spacer sets and the position of TRE units in the DNA helix relative to the minimal promoters for specific TF motifs (Fig. S4B-D).

**Figure 2.**
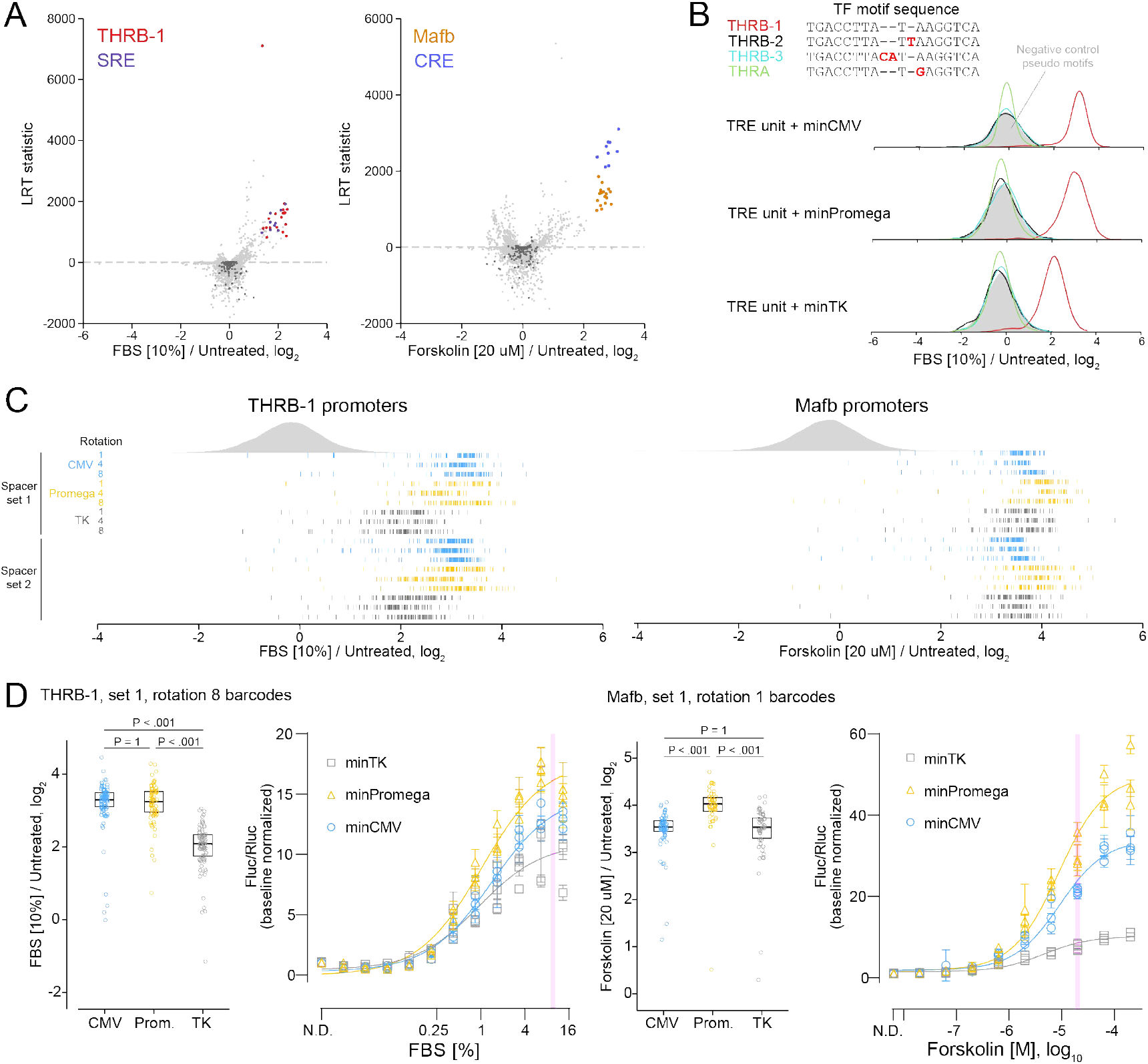
TRE-MPRA benchmarking with fetal bovine serum and forskolin. **A)** Volcano plots of promoter responses following stimulation of HEK293 cells with 10% FBS (left) or 20 uM forskolin (right) for six hours, in comparison to untreated cells. Select promoters of interest are colored by TFBM. Dashed lines indicate an FDR threshold of 5%. Negative control promoters are indicated by dark gray data points. LRT, likelihood ratio lest. **B**) Density plots of the barcode fold changes for promoters containing the THRB-1 motif or motifs of similar sequence following stimulation with 10% FBS. Other motifs with similar sequence to THRB-1 showed no effect upon FBS treatment. **C**) Raincloud plots of fold changes for individual barcodes across each THRB-1 (left) or Mafb (right) promoter in FBS or forskolin treated HEK293 cells, respectively, relative to untreated controls. Each barcode is represented by a vertical line. Density plots indicate negative control TRE barcodes. **D**) THRB-1 and Mafb TRE promoter responses tested in orthogonal dual-luciferase assays. Boxplots show fold changes for individual barcodes of the THRB-1 unit with spacer set 1 and a span of eight (left) or the Mafb unit with spacer set 1 and a span of 1 (right) paired with each minimal promoter from the TRE-MPRA experiment. Boxes indicate the median and interquartile range. P-values indicated are from one-way analysis of variance tests with Sidak adjustments for multiple comparisons. Associated dose-response curves of the same TRE units obtained from dual-luciferase assays. Data were scaled (‘baseline normalized’) to the Fluc/Rluc ratio in untreated cells (‘N.D.’). Curves were fit to scaled Fluc/Rluc values across three experimental replicates. Data points and error bars indicate the mean and standard deviation of four technical replicates within each experimental replicate. Shaded vertical lines indicate the drug doses used in the TRE-MPRA experiment.

For each barcode, we calculated the median reads per million across biological replicates and then compared barcode fold changes (treatment versus unstimulated) of all THRB-1 and Mafb promoters against the panel of negative control promoters (Fig. 2C). Consistent responses across the populations of barcodes suggested the induction of individual promoters following stimulation was highly reproducible. We also noted the degree of induction of THRB-1 and Mafb TRE activity was dependent upon the paired minimal promoter (Fig. 2D). To test these findings in an orthogonal assay, we derived dual-luciferase reporter plasmids containing individual promoters controlling the expression of a luc2P CDS, as well as a constitutive SV40-driven Renilla luciferase cassette. HEK293 cells transfected with reporter plasmids showed significant, dose-dependent increases in relative luc2P activity following stimulation with FBS or forskolin (Fig. 2D). Furthermore, minimal promoter-dependent responses were observed, largely in agreement with our TRE-MPRA results (Fig. 2D). These results demonstrate that our synthetic promoters can function as dynamic transcriptional readouts of cell signaling.

These orthogonal dual-luciferase experiments also found baseline transcription rates of the THRB-1 and Mafb promoters in untreated cells to be in agreement with the estimated transcription rates in untreated cells generated from the MPRA platform (Fig. S5A). To further determine if constitutive transcription rate estimates derived from our TRE-MPRA library are reliable predictors of expression output in orthogonal assays, we derived and tested dual-luciferase reporter plasmids containing promoters spanning a range of estimated transcription rates from untreated HEK293 cells (Fig. S5B). This series of reporters produced luciferase activities in line with the MPRA transcription rate estimates, with the exception of the BHLHB3 motif-containing promoter (Fig. S5C). This discordant finding may reflect TF/TRE-specific translation rates observed in previous studies^22–24^, and emphasizes the necessity to validate individual candidate promoters in orthogonal assays.

### Identifying synthetic promoters responsive to endogenous cellular responses

With a validated MPRA platform in hand, we next sought to modulate the activities of additional promoters in the library by treating HEK293 cells with eight additional stimuli, including mitogens and inducers of cellular stress (Table S1). To increase throughput, we included a single replicate for most treatment conditions, as our preliminary results from FBS and forskolin treated cells showed that individual biological replicates identified the same sets of top responding promoters as multiple replicates (Fig. S6). We performed hierarchical clustering to classify common and unique promoter responses (Fig. 3A). For most stimulus types, we observed hundreds of promoters with significantly altered activity, even at a stringent FDR cutoff of 0.05%, attesting to the statistical power of MPRAs (Fig. 3B). Across ten stimulus conditions, 4342 promoters (70.7%) showed altered transcriptional output in at least one condition relative to negative controls, while 824 promoters (13.4%) were altered in at least half of the conditions. Several treatments activated promoters with similar or greater effect sizes as were seen for FBS- or forskolin-responsive promoters. For example, dexamethasone treatment caused significant (2-6 fold) upregulation of TRE units containing motifs for two nuclear receptor superfamily class I (steroid) members: androgen receptor (AR) and glucocorticoid receptor (NR3C1) (Fig. 3C). A dual-luciferase plasmid containing an AR promoter showed a dose-dependent response to dexamethasone with a dynamic range of 134-fold (Fig. 3D). In addition, treatment with lithium chloride caused strong activation of TRE units with an NFAT5 motif (Fig. 3E), in line with lithium’s inhibition of glycogen synthase kinase 3 beta, itself an inhibitor of NFAT5 transcriptional activity^25–28^.

**Figure 3.**
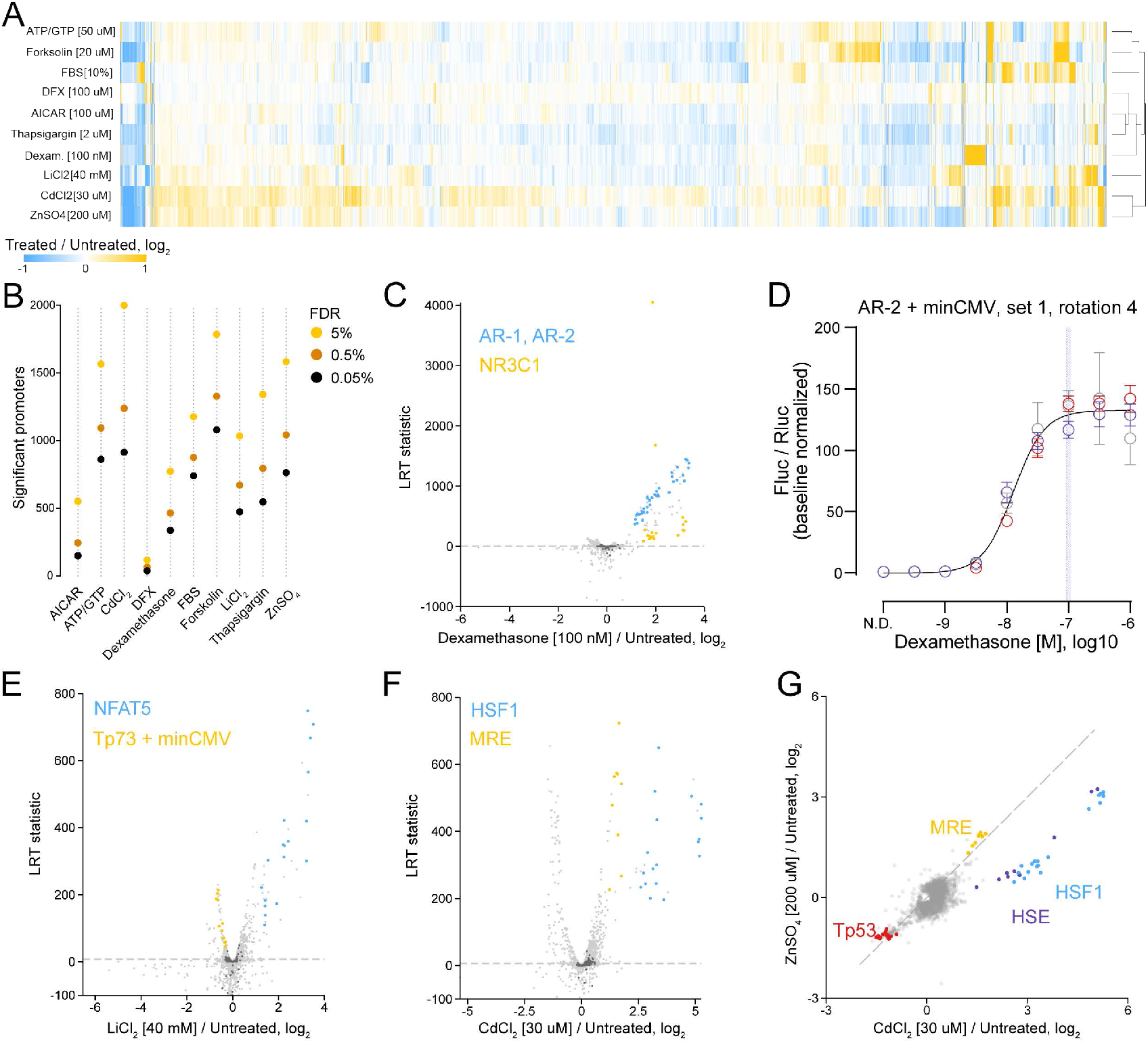
TRE-MPRA results using additional stimuli. **A**) Heatmap of protein fold change responses in HEK293 cells across ten stimulus conditions relative to untreated cells. Treatment conditions were clustered using Euclidean distance with complete linkage. **B**) The number of significantly altered promoters upon stimulation across varying FDR cutoffs. **C**) Volcano plot of promoter responses following treatment of HEK293 cells with 100 nM dexamethasone, 40 mM lithium chloride, or 30 uM cadmium chloride for six hours, in comparison to untreated cells. Dashed lines indicate an FDR threshold of 5%. Negative control promoters arc indicated by dark gray data points. **D**) Dose response curves from HEK293 cells transfected with an AR promoter dual luciferase reporter and treated with dexamethasone. Data were scaled (‘baseline normalized’) to the Fluc/Rluc ratio in untreated cells (‘N.D.’). The curve was fit to baseline normalized Fluc/Rluc values across three experimental replicates. Data points and error bars indicate the mean and standard deviation of four or eight technical replicates (n=4,8) within each of three experimental replicates (N=3, distinguished via color). The shaded vertical line indicates the dose of dexamethasone used in the TRE-MPRA experiment. **E**,**F**) Volcano plot of promoter responses following treatment of HEK293 cells with 40 mM lithium chloride (E) or 30 uM cadmium chloride (F) for six hours, in comparison to untreated cells. Dashed lines indicate an FDR threshold of 5%. Negative control promoters are indicated by dark gray data points. **G**) Scatterplot comparing promoter responses between lithium and cadmium treatments. HSF1 units were more responsive to cadmium. Dashed line indicates the identity line (y=x).

Our treatment conditions included two heavy metals: one physiologic (zinc) and one xenobiotic (cadmium). We noted that both metal treatments induced similar responses in metal response element (MRE)-containing and Tp53-containing promoters (Fig. 3F,G). The MRE is bound by metal regulatory transcription factor 1 (MTF-1), in response to both zinc and cadmium elevation^29^, as well as oxidative stress^30^. However, cadmium treatment showed two- to three-fold higher induction of HSF1- and heat shock element (HSE)-containing promoters relative to zinc. Previously, using a modified HEK293 cell line containing a HSE reporter^31^, it was observed that HSE was roughly 200-fold more sensitive towards cadmium than zinc^32^. Because HSE is bound by both MTF-1 and HSF1, the observed HSF1 TRE activation discrepancy between zinc and cadmium treatments is likely due to HSF1 induction rather than MTF-1^33^.

We then transfected the TRE-MPRA library into mouse Neuro2a cells in order to compare baseline transcription rates between cell lines from different species and tissue origins. Promoters were classified as being transcriptionally active or inactive in each HEK293 experimental replicate and in Neuro2a cells using MPRAnalyze (Fig. S7A). We observed near perfect agreement in promoter classification between HEK293 experimental replicates (Fig. S7B,C). Sets of active and inactive promoters were also significantly enriched between HEK293 and Neuro2a cells, but the overlaps were much less pronounced. Furthermore, the estimated transcription rates for overlapping sets of active promoters between HEK293 and Neuro2a cells showed much lower correlations compared to the HEK293 experimental replicate comparison (Fig. S7C). Next, we treated Neuro2a cells with 10% FBS for six hours to profile promoters responding to serum and compared those to responding promoters in HEK293 cells. The TRE units with the strongest response to FBS in Neuro2a cells contained the MRE motif, while the strongest responding promoters in HEK293 cells were much less responsive in Neuro2a cells (Fig. S7D,E). Because baseline promoter transcription rates and the responses to serum were discordant between the two cell lines tested, we recommend performing the TRE-MPRA screen in specific cell models of interest to identify optimal elements.

### Specific synthetic promoters activated by aminergic GPCR agonism

After identifying synthetic promoters with large dynamic range responses across multiple chemical and mitogen treatments, we next examined whether the TRE-MPRA library possesses the sensitivity necessary to detect transcriptional responses following GPCR activation. A limited number of TREs have long been used as readouts for GPCR signaling events, particularly as part of bioluminescent sensors^34^. However, the specificity of existing TREs as readouts for a particular GPCR-G protein coupling is not well supported experimentally; furthermore, the complexity of GPCR signaling suggests paring down receptor activation to a single, G protein-specific TRE is unlikely^35–37^. Here, we employed TRE-MPRA to profile transcriptional changes induced by activation of three well-characterized aminergic GPCRs. HEK293 cells were co-transfected with the TRE-MPRA plasmid library and a plasmid expressing one of three human GPCRs: β_2_ adrenergic receptor (ADRB2), 5-hydroxytryptamine receptor 2A (HTR2A), or dopamine receptor D2 (DRD2). These receptors selectively couple to Gα_s_-, Gα_q_-, and Gα_i_-containing heterotrimeric G protein complexes, respectively. Upon GPCR activation by receptor agonists, these G protein complexes initiate distinct downstream signaling cascades^38,39^. After six hours of receptor agonist treatment (ADRB2: 1 uM epinephrine; HTR2A: 100 nM 5-HT; DRD2; 1 uM dopamine), we harvested RNA and analyzed differential barcode abundance between receptor alone and receptor with agonist conditions (Fig. 4A). Epinephrine triggered increased expression from promoters containing Mafb and CRE motifs, similar to what we observed in forskolin treated cells, as both forskolin and Gα_s_ signaling stimulate the formation of cyclic AMP. Conversely, dopamine binding to DRD2 activates Gα_i_ signaling, which leads to the inhibition of cyclic AMP production^40–42^. As such, we saw distinct promoters become active upon DRD2 agonism. Notably, effect sizes for promoters significantly responding to dopamine treatment were substantially smaller than those observed for the other two receptor agonists. To identify promoters that selectively distinguish between Gα_s_, Gα_q_, and Gα_i_ signaling cascades, we generated biplot displays for the set of promoters that showed a significant response to receptor agonism in at least one of three GPCR comparisons (Fig. 4C)^43^. Projections of promoter fold changes segregated those promoters responding uniquely to GPCR agonism.

**Figure 4.**
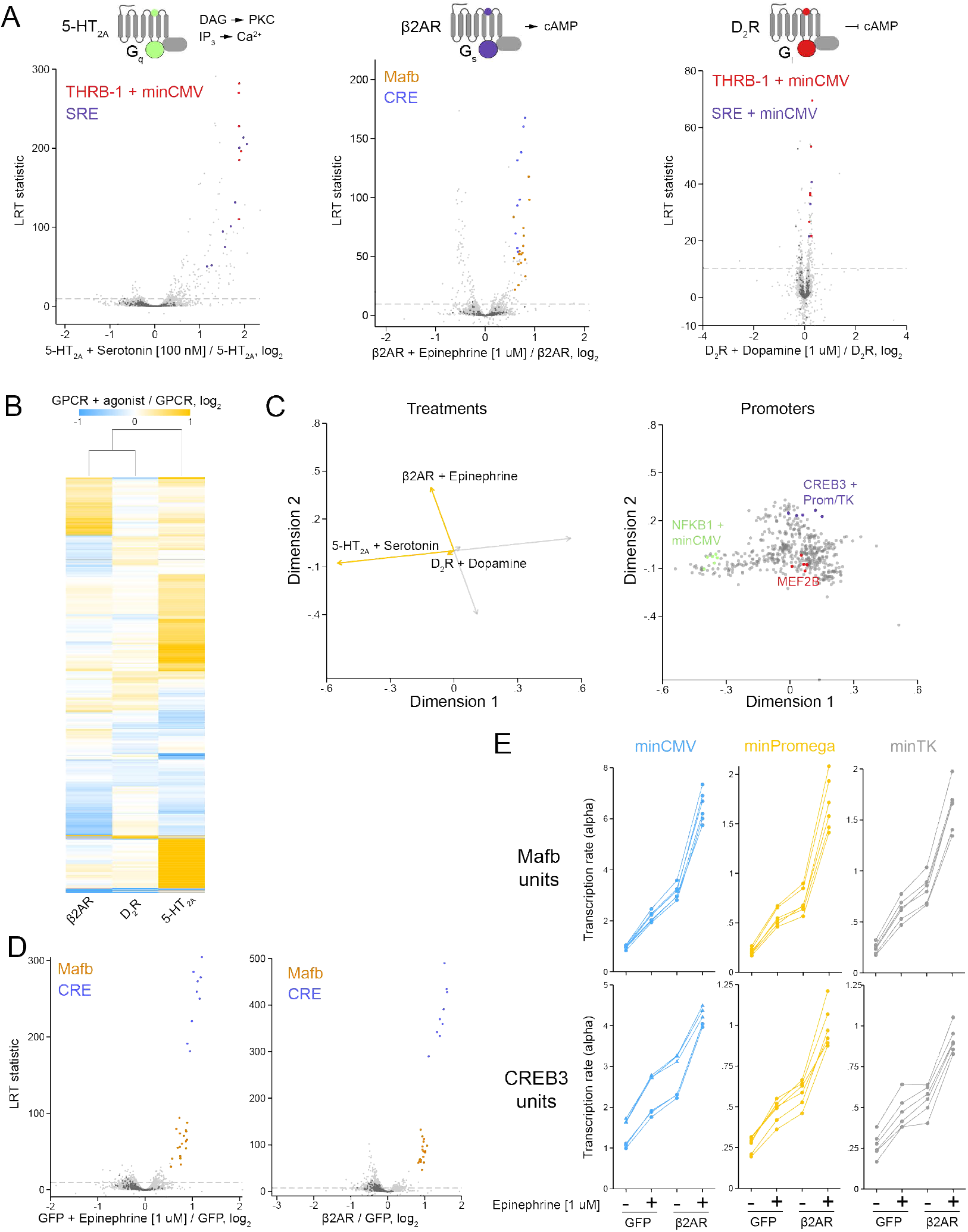
TRE-MPRA detects distinct transcriptional signals downstream of aminergic GPCR agonism. **A)** Volcano plots of promoter responses following receptor agonism in HEK293 cells co-transfected with the TRE-MPRA library and GPCR expression plasmids. Dashed lines indicate an FDR threshold of 5%. Negative control promoters arc indicated by dark gray data points. **B)** Hcatmap of promoter fold change responses to aminergic GPCR agonism in HEK293 cells. Displayed are the fold change values for all promoters that were significantly altered by agonist treatment in one comparison.**C)** Separated biplots displaying the treatment (left) and promoter (right) projections across two dimensions. GPCR-specific responding promoters arc highlighted. Orange and gray projection lines indicate the positive and negative directions of the treatment projection, respectively. **D)** Volcano plots comparing promoter responses of HEK.293 cells treated with 1 uM epinephrine and untreated cells (left) and HEK293 cells transfected with GFP or ADRB2 expression plasmids (right).Dashed lines indicate an FDR threshold of 5%. Negative control promoters arc indicated by dark gray data points. **E)** Transcription rate estimations (alphas) for Mafb and CREB3 units in HEK293 cells following 1 uM epinephrine treatment or ADRB2 overexpression, or both. Each connected set of data points represents a single promoter. In the min CMV CREB3comparison, the filled triangular data points represent units with spacer set 2, filled circular points arc units containing spacer set 1.

Having detected transcriptional changes mediated by exogenous, overexpressed GPCR agonism, we next asked whether the TRE-MPRA platform can detect transcriptional changes due to endogenous GPCR agonism, as well as changes resulting from receptor overexpression in the absence of an exogenous ligand. To address this question, we focused on ADRB2 in HEK293 cells, which are known to express functional ADRB2 endogenously^44^. We co-transfected cells with the TRE-MPRA library and a control plasmid expressing GFP and then cultured the cells in serum-free media in either the presence or absence of 1 uM epinephrine for six hours. Epinephrine treatment alone resulted in elevated transcription from promoters that had responded to epinephrine treatment in the presence of overexpressed ADRB2, suggesting that this platform is sensitive enough to detect endogenous GPCR signaling following agonism (Fig. 4D, left). Furthermore, the overexpression of ADRB2 in the absence of epinephrine treatment also caused increased transcription from these promoters, suggesting that either basal constitutive signaling from, or autocrine activation of, plasmid-expressed GPCRs can be detected with our platform (Fig. 4D, right). Estimated transcription rates across the four treatment conditions for promoters containing Mafb or CREB3 units were sensitive to endogenous and exogenous ADRB2 signaling to similar degrees across the three minimal promoter combinations (Fig. 4E). Notably, the spacer set effect on CREB3-minCMV promoters seen following forskolin treatment (Fig. S4B) was also observed in these data (Fig. 4E).

### Synthetic promoter activation following agonism of promiscuously coupled non-aminergic GPCRs

Having profiled synthetic promoter activity following agonist treatment of canonical Gα_s_-(ADRB2), Gα_q_-(5-HT2A), and Gα_i_-coupled (DRD2) aminergic GPCRs and identified G protein-specific changes following activation, we next used the TRE-MPRA platform to profile activation of three additional GPCRs: the adhesion class protease-activated receptor-1 (PAR1), the recently de-orphaned succinate receptor (GPR91)^45^, and the MAS-related G Protein-Coupled Receptor-X2 (MRGPRX2)^46–51^, which can strongly couple to and activate multiple distinct G proteins in HEK293 cells^52^. For each of these GPCRs, agonist treatment significantly upregulated many promoters that were also increased by HTR2A agonism, such as THRB-1 and SRE-containing constructs, suggesting that these GPCRs induce transcriptional changes predominantly via Gα_q_ signaling (Fig. 5A). Indeed, PAR1, GPR91, and MRGPRX2 profiles showed higher correlation with HTR2A than with ADRB2 or DRD2 (Fig. 5B). To more thoroughly compare the six GPCR profiles, we again generated biplot displays using the set of promoters that showed a significant response to receptor agonism in at least one of the six datasets (Fig.5C). Projections for PAR1, GPR91, and MRGPRX2 conditions were closely related to that of HTR2A, again suggesting these GPCRs may activate similar downstream signaling pathways. However, these three projections were most similar to each other. Among promoters most responsible for the distinction from HTR2A were those containing MEF2B and FOXB1 units, which showed higher activation in PAR1, GPR91, and MRGPRX2 conditions compared to HTR2A (Fig. 5C,D).

**Figure 5.**
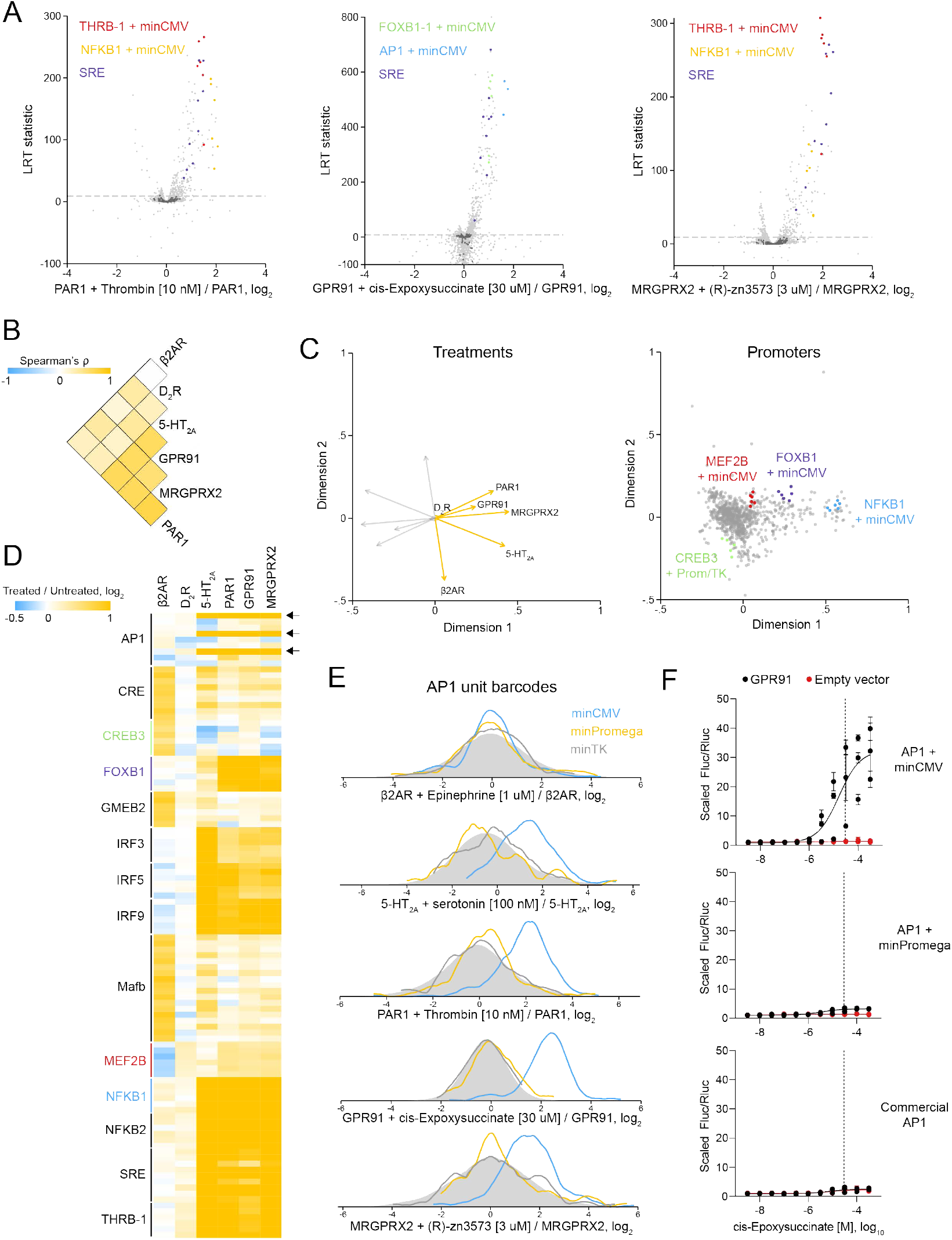
Profiling promoter activity following activation of additional GPCRs. **A)** Volcano plots of promoter responses in HEK293 cells co-transfectcd with the TRE-MPRA library and GPCR expression plasmids following receptor agonism. Dashed lines indicate an FDR threshold of 5%. Negative control promoters arc indicated by dark gray data points. **B)** Hcatmap of Spearman’s correlation coefficients for all promoters significantly altered by agonist treatment in at least one of six GPCR experiment groups. **C)** Separated biplots for the set of promoters that showed a significant response to receptor agonism in at least one of the six datasets. Displayed arc the treatment (left) and promoter (right) projections across two dimensions. Responding promoters of interest arc highlighted. Orange and gray projection lines indicate the positive and negative directions of the treatment projection, respectively. **D)** Hcatmap of selected promoter responses across GPCR experiment groups. Arrows indicate the API unit/minCMV promoters. A list of promoter (row) labels can be found in TableS6. **E)** Density plots of barcode fold changes following receptor agonism for API units by minimal promoter. The filled gray distributions indicate the population of negative control promoters. **F)** Dose response curves from HEK.293 cells co-transfected with API promoter (rotation of four) dual luciferase reporters and either an empty plasmid or a GPR91expression plasmid. Data were sealed to the Fluc/Rluc ratio in untreated cells (set to a value of 1). Curves were fit to scaled Fluc/Rluc values across three experimental replicates. Data points and error bars indicate the mean and standard deviation of four technical replicates within each experimental replicate. Vertical dashed lines indicate the drug dose used in the TRE-MPRA experiment.

We also noted that activation of AP1 units by HTR2A, PAR1, GPR91, and MRGPRX2 agonism was strictly dependent on the minCMV promoter pairing (Fig. 5D,E). To validate this striking finding, we generated dual-luciferase reporters that replicated the sequences of minCMV and minPromega versions of AP1 TRE units of the MPRA library. We also converted a commercially available AP1 luciferase reporter into a dual-luciferase reporter by replacing its hygromycin expression cassette with an SV40/Rluc cassette for direct comparison with our synthetic promoters. We then co-transfected HEK293 cells with a GPR91 expression plasmid and the AP1 reporters and measured firefly and Renilla luciferase activities after six hours of cis-epoxysuccinate treatment. We saw a dose-dependent increase in transcriptional output for the AP1 unit coupled with the minCMV, whereas the minPromega coupling and the modified commercial AP1 reporter showed little or no response to cis-epoxysuccinate, in agreement with the results of the TRE-MPRA experiment (Fig. 5F).

## Discussion

Incorporating promoters with desired response profiles in reporter constructs enables precise monitoring of cell signaling and the development of synthetic biology applications where programmable transcription readouts are essential. This study introduces a novel MPRA library designed to simultaneously screen thousands of candidate synthetic promoters in mammalian cells. Using this platform, we identified promoters active in diverse cell lines and under various stimuli. This system will streamline the development of complex gene regulatory networks and offers a valuable tool for the rational design of synthetic gene circuits with stringent input-output relationships.

We orthogonally tested several synthetic promoters in dual-luciferase assays and observed dose-dependent responses to stimulus, with dynamic ranges of 15- to 134-fold relative to baseline. Because our screening library contains only a handful of configurations for each candidate TRE, and all promoters were built with homotypic TREs, there is likely considerable room for optimization to further enhance performance upon orthogonal validation. Indeed, heterotypic TRE combinations with elevated transcriptional activity relative to homotypic promoters have been reported^53^.

The MPRA format enables the simultaneous measurement of multiple technical replicates of the same genetic sequence of interest, which provides a high level of statistical power to detect even minor effect sizes. Our experiments consistently detected promoters with effect sizes of less than 15% between treatment groups while maintaining false discovery rates below 5%. Therefore, the TRE-MPRA library can effectively capture changes in promoter activity that might be missed by other screening formats. It’s worth noting that the most suitable promoter for a given application might not be one of the largest responders in a specific screen. For example, in applications where a range of transgenic expression levels is desired to assess concentration-effect sufficiency.

As a demonstration of the TRE-MPRA library’s sensitivity, we were able to detect changes in promoter activity as a result of the agonism of endogenous ADRB2 in HEK293 cells. Furthermore, the overexpression of ADRB2 in the absence of ligand activated these same promoters. We anticipate our TRE-MPRA library will have broad applications for studying GPCR signaling. GPCRs have traditionally been associated with activating Gα, β, γ transducers and arrestins; however, recent research has revealed unique signaling properties beyond these paradigms, including coupling to non-G-protein elements or inducing cellular signaling from endosomal compartments^54–59^. By profiling promoter activity changes in response to GPCR activation users can obtain an unbiased view of total cellular signal transduction that can be converted directly into reporters, even for pathways of unknown molecular origin.

## Materials and Methods

### Reagents and drug preparation

Fetal bovine serum was purchased from Omega Scientific. Cis-Epoxysuccinic acid was purchased from ThermoFisher Scientific. Forskolin, dopamine, thapsigargin, and (R)-zn3573 were purchased from Tocris Bioscience. ATP and GTP were purchased from New England Biolabs. All other chemicals were purchased from Sigma-Aldrich. Dilutions of stock solutions were made in 3X drug assay buffer (0.3 mg/mL ascorbic acid, 0.3% bovine serum albumin, and 20 mM HEPES in HBSS).

### Transcription factor binding motif selection

TF binding motifs included in the initial screening library were selected from a published set of position weight matrices (PWMs) derived for 411 human and mouse TFs using HT-SELEX^17^. Beginning with the seed sequence of each PWM, we first trimmed fully degenerate nucleotides (Ns) from the 5’ and 3’ ends and then replaced all remaining degenerate nucleotides with the predominant nucleotides at those positions. Finally, to eliminate redundancy in our candidate list, we removed any resulting motif for which the entire sequence was represented within another motif, resulting in a total of 325 unique binding motifs (Table S2).

### TRE unit design

For each experimental binding motif, we designed six unique TRE units, each consisting of four copies of the binding motif separated by random nucleotide spacer sequences, a random sequence of variable length following the 3’ motif, and flanked by restriction enzyme recognition sequences and primer binding sites (Fig S1). We chose to test homotypic TRE units containing four copies of the binding motifs based on a previous report showing maximal activity with synthetic promoters containing four copies of a cAMP response element^60^. Spacer lengths were selected based on individual motif lengths such that adjacent motifs were oriented on opposite sides of the DNA double helix. To the resulting list of 1950 TRE units, we added 54 TRE units based on Promega’s pGL4 Luciferase Reporter Vectors ranging in length from 160 to 194 nucleotides, and 102 negative control TRE units, for a total of 2106 TRE units (Table S3). Each TRE unit was examined for compatibility with our restriction enzyme strategy and modified as necessary to remove unwanted recognition sites. All TRE unit oligonucleotides, except those encoding positive control TREs greater than 160 nucleotides, were synthesized as 160 nucleotides in length by adjusting sequence lengths 3’ to the TRE unit to limit bias in PCR amplification prior to library assembly.

### Barcoding of TRE units

Synthesized oligonucleotides encoding TRE units were purchased from Twist Bioscience and pre-amplified via qPCR (Fig. S1A, Table S4). Following pre-amplification, barcodes were added to the 3’ end of the amplicons using degenerate primers during 10 cycles of qPCR in six distinct reactions. Each reaction was processed separately for the remainder of the library generation procedure. Amplicons were digested with MluI and SpeI and ligated to pDonor_eGP2AP_RC (Addgene #133784) digested with MluI and SpeI in triplicate reactions. Each ligation reaction was transformed separately into NEB 10-beta competent cells in triplicate. Following transformation, triplicates were pooled and cultured in 2X YT media containing 50 ug/mL kanamycin at 30 degrees C with shaking at 225 rpm overnight.

To generate TRE unit/barcode dictionaries, dual-indexed amplicons were generated from the purified plasmid pools via PCR. We generated and sequenced two amplicons from each replicate using unique indices. Amplicons were sequenced on a NextSeq 550 (Illumina) using a NextSeq 500/550 High Output 300 cycle flow cell with custom sequencing primers (Table SX). TRE unit and barcode sequences were extracted from demultiplexed read pairs. Resulting TRE unit sequences were compared to the expected sequences using Starcode^61^. TRE unit/barcode pairs for which the TRE unit was within a Levenshtein distance of two from the expected unit sequence were retained. Any barcode that was paired with multiple unique TRE units was discarded from the dictionary.

### TRE-MPRA plasmid library assembly

After generating the TRE unit/barcode dictionaries, fragments containing a firefly luciferase CDS and one of three minimal promoters – thymidine kinase (minTK), the minimal promoter of Promega’s pGL4 plasmid suite (minProm), or cytomegalovirus (minCMV) – were created in a modified pDonor_eGP2AP_RC via Gibson Assembly, digested with KpnI and XbaI, and then ligated into the barcoded plasmid library replicates using the KpnI and XbaI restriction enzyme sites. Each replicate received one of the three minimal promoters.

Plasmid libraries for transfection were prepared by inoculating 100 mL of 2X YT media containing 50 ug/mL kanamycin with 200 uL of bacterial glycerol stocks and incubating at 30 degrees C with shaking at 225 rpm overnight. Cultures were pelleted and plasmids were purified using a Plasmid Maxiprep Kit (Qiagen) according to manufacturer’s protocol. Plasmid preparations were then combined at equimolar ratios. TRE-MPRA libraries are available from Addgene under deposit number 82594.

### Cell culture and TRE-MPRA plasmid library transfection

HEK293 cells were maintained in DMEM (4.5 g/L D-glucose) supplemented with 10% fetal bovine serum, penicillin (100 IU/mL) and streptomycin (100 ug/mL) (‘growth media’). Neuro2a cells were maintained in EMEM supplemented with 10% fetal bovine serum, penicillin (100 IU/mL) and streptomycin (100 ug/mL) (‘growth media’).

For experiments with the full TRE-MPRA library, 5 × 10^6^ cells were plated on 15 cm treated tissue culture dishes in growth media. The following day, cells were transfected with 10 ug of the TRE library ± 5 ug of pcDNA3.1 containing a GPCR expression cassette using TransIT-2020 (Mirus Bio) according to manufacturer’s instructions. After 6 hours, cells were washed with serum-free (SF) versions of growth media (‘SF media’) and then cultured overnight in SF media. The following day, stimulus was added to the media and cells were cultured for an additional 6 hours. Cells were then trypsinized using 0.05% trypsin with EDTA, pelleted by centrifugation, and frozen at -80 degrees C until processing. Treatment conditions are listed in Table S1.

### Nucleic acid isolation

RNA fractions from cell pellets were isolated using QIAshredder homogenizers and the AllPrep DNA/RNA Mini Kit (Qiagen). Immediately after homogenization, a pool of four synthesized spike-in RNAs (2.5 fM each, 10 fM total) was added to each 600 uL sample (Table S4). Spike-in RNAs served to identify any samples with poor barcode recovery. RNA fractions were eluted in 30 uL nuclease-free water. Following isolation, the RNA fraction was treated with TURBO DNase (Invitrogen) to remove any plasmid DNA carryover. DNA removal from RNA fractions was confirmed by RT-PCR using the SuperScript IV One-Step RT-PCR System (Invitrogen) with and without addition of SuperScript IV RT Mix, followed by agarose gel electrophoresis. RT-PCR was performed according to manufacturer’s instructions with the following conditions: 55 C for 10 minutes, 98 C for 2 minutes, 30 cycles of PCR (98 C for 10 seconds, 60 C for 10 seconds, 72 for 8 seconds), and 72 C for 5 minutes. Primer sequences are listed in Table S4.

### Sequencing library generation

Dual-indexed amplicons from RNA samples were generated using the SuperScript IV One-Step RT-PCR System (Invitrogen). 0.5-1.0 uL of RNA was used as the template in 20 uL reactions. RT-PCR was performed according to manufacturer’s instructions with the following conditions: 55 C for 10 minutes, 98 C for 2 minutes, 17-27 cycles of PCR (98 C for 10 seconds, 60 C for 10 seconds, 72 for 8 seconds), and 72 C for 5 minutes. Dual-indexed amplicons from plasmid DNA samples were generated using the Platinum SuperFI II Green PCR Master Mix (Invitrogen). PCR was performed according to manufacturer’s instructions with the following conditions: 98 C for 30 seconds, 16 cycles of PCR (98 C for 5 seconds, 60 C for 10 seconds, 72 C for 5 seconds), and 72 C for 5 minutes. Products were separated by agarose gel electrophoresis and library amplicons were extracted using the QIAquick Gel Extraction Kit (Qiagen). Amplicons were quantified with the KAPA Library Quantification Kit for Illumina Platforms (KAPA Biosystems) in 384-well format using a CFX Opus 384-Well Real-Time System (Bio-Rad). Libraries prepared from RNA samples were pooled at equimolar concentrations and combined with plasmid DNA input amplicons such that plasmid amplicons represented approximately 6-8% of the pool. RNA and plasmid DNA libraries were sequenced using a 50 cycle SP flow cell on a NovaSeq 6000 (Illumina) using custom sequencing primers (Table S4).

### Processing of sequencing data

Our processing pipeline was built using Snakemake^62^. Raw barcode counts from TRE-MPRA samples were derived from demultiplexed fastq files by collecting the first 24 nucleotides of each read. Paired-end reads were trimmed using TrimGalore^63^ and joined using FastQ-Join^64,65^. Candidate barcodes were clustered within each sample via Starcode using a Levenshtein distance of one^61^. Reads of each barcode cluster were then cross-referenced with the barcode dictionaries and only the candidate clusters with either 1) the centroid present in the dictionaries, and all other collapsed reads not present, or 2) the centroid present and any other collapsed reads present in the dictionary mapping to the same synthetic promoter, were retained for analysis. Reads within each retained clusters were collapsed into a sum total for the cluster and assigned to the centroid barcode. Reads mapping to RNA spike-in sequences were also tallied within each sample. Spike-in reads were used to assess individual sample quality by flagging samples with disproportionately low proportions of promoter barcode reads; no such samples were observed in our datasets.

### Expression plasmid derivation

Plasmids used in this study are listed in Table S5. pcDNA3.1_eGFP was generated by PCR amplifying the eGFP CDS from Arch(D95H)-eGFP (Addgene #51081) and inserting into pcDNA3.1(-)/*myc*-His A using the EcoRI and NotI restriction sites. pcDNA3.1_Signal-Flag-ADRB2 was generated by PCR amplifying the ADRB2 CDS from ADRB2-Tango (Addgene #66220) (introducing a stop codon) and inserting into pcDNA3.1(-)/*myc*-His A using Gibson assembly. pcDNA3.1_Signal-Flag-DRD2 was generated by PCR amplifying the DRD2 CDS from DRD2-Tango (Addgene #66269) (introducing a stop codon) and inserting into pcDNA3.1(+) using the NotI and XhoI restriction sites. pcDNA3.1_Signal-Flag-HTR2A was generated by PCR amplifying the HTR2A CDS from HTR2A-Tango (Addgene #66409) (introducing a stop codon) and inserting into pcDNA3.1(+) using the NotI and XhoI restriction sites. pcDNA3.1-GPR91 was generated by PCR amplifying the GPR91 CDS from SUCNR1-Tango (Addgene #66507) (introducing a stop codon) and inserting into pcDNA3.1(-)/*myc*-His A using Gibson assembly. pTwist-PAR1 was purchased from Twist Bioscience. pcDNA3.1_Signal-Flag-MRGPRX2 was previously described^66^.

### Dual-luciferase TRE reporter plasmid derivation

The hygromycin CDS of pGL4.33[*luc2P*/SRE/Hygro] (Promega) was replaced with a Renilla luciferase CDS via Gibson assembly using NEBuilder HiFi DNA Assembly Master Mix (New England Biolabs) to generate a dual-luciferase reporter containing luc2P and Renilla expression cassettes (pGL4.33R). To generate a TRE unit acceptor site upstream of the minimal Promega promoter replicating the promoter sequence of our TRE-minPro screening plasmids, annealed oligos containing restriction enzyme recognition sites were ligated to pGL4.33R sequentially digested with BglII and KpnI (pGL4.R_TRE_minPro). To derive a minimal CMV promoter/TRE driven dual-luciferase plasmid, the minimal CMV promoter region of the TRE-MPRA minCMV plasmid was first PCR amplified, digested with ApaI and KpnI, and ligated to pGL4.33R digested with ApaI and KpnI (pGL4.R_minCMV). Next, to replicate the promoter sequence of our TRE-minCMV screening plasmids, annealed oligonucleotides containing restriction enzyme recognition sites were ligated to pGL4.R_minCMV digested with KpnI (pGL4.R_TRE_minCMV). To derive a minimal TK promoter/TRE driven dual-luciferase plasmid, the TK promoter fragment of the TRE-minTK library was ligated to pGL4.R_minCMV digested with ApaI and KpnI (pGL4.R_minTK). Next, to replicate the promoter sequence of our TRE-minTK screening plasmids, annealed oligonucleotides containing restriction enzyme recognition sites were ligated to pGL4.R_minTK digested with KpnI (pGL4.R_TRE_minTK).

Annealed oligonucleotides corresponding to individual TRE units were ligated into pGL4.R_TRE plasmids digested with MluI and KpnI (minCMV and minPro) or AscI and KpnI (minTK) to generate dual-luciferase reporters with synthetic promoters identical to those of the MPRA constructs controlling expression of the luc2P CDS.

### Dual-luciferase assay

HEK293 cells were plated on 10 cm tissue culture-treated dishes in growth media. The following day, cells were transfected with 1 ug of TRE reporter plasmid using TransIT-2020 according to manufacturer’s instructions. After 6 hours, cells were detached from culture dishes with 0.05% trypsin, washed in SF media and plated on poly-d-lysine coated 384-well plates (10,000 cells/well in 20 uL of SF media). The following day, 10 uL of drug dilutions in SF media were added to wells and plates were incubated for six hours. Firefly and Renilla luciferase activities were then measured sequentially on a Pherastar using the Dual-Glo Luciferase Assay (Promega, catalog #E2920) according to manufacturer’s instructions. For GPR91 experiments, HEK293 cells were plated on 10 cm tissue culture-treated dishes in growth media and co-transfected the following day with 1 ug of TRE reporter plasmid and 3μg receptor plasmid using TransIT-2020 according to manufacturer’s instructions. After 6 hours, a media change was performed to replace growth media with 1% d-FBS media (DMEM with 4.5 g/L D-glucose supplemented with 1% dialyzed fetal bovine serum, penicillin 100 IU/mL and streptomycin 100 ug/mL) and cells were incubated overnight. The next day, cells were detached from culture dishes with Dulbecco’s phosphate-buffered saline containing 500 uM EDTA and plated on poly-d-lysine coated 384-well plates (10,000 cells/well in 20 uL of 1% d-FBS media). The following day, 10 uL of drug dilutions in drug assay buffer^37^ were added to wells and plates were incubated for six hours. Firefly and Renilla luciferase activities were then measured sequentially as above.

### Transcription rate estimation and comparative analyses

Estimations of promoter transcription rates and comparisons between treatments were performed using MPRAnalyze^18^. MPRAnalyze utilizes raw NGS read data of individual barcodes to generate a single transcription rate estimate for each promoter using generalized linear models. Similarly, raw barcode data is used to perform comparisons of individual promoters between treatment groups. For each promoter with more than 100 associated barcodes, we selected the 100 barcodes with highest abundance in the plasmid DNA libraries for inclusion in these analyses to reduce computation overhead. All other promoters had all barcodes included in the analyses. Individual sample read depth factors were calculated by scaling the upper quartile of raw read counts to that of an arbitrarily chosen reference sample (upper quantile read counts/ reference sample upper quartile read counts).

Within a treatment condition, individual promoter transcription rates were compared to the rates of negative control promoters using median absolute deviation (MAD) scores from the ‘testEmpircal’ command of MPRAnalyze. MAD score p-values were then corrected for multiple testing using the step-up method of Simes in the ‘qqvalue’ Stata package^67^. Promoters with resulting FDR values less than 0.05 (5%) were considered transcriptionally active in the condition. The fitted models for transcription rate estimations were:

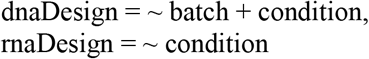

To compare activity between treatment conditions, MPRAnalyze performs likelihood ratio tests (LRTs). Because many resulting p-values and false discovery rates were less than 10^−38^ and were thus outputted as zeros, we have chosen to display the resulting LRT statistic in our volcano plots. For those interested, the reported LRT statistics can be converted to p-values using a chi-square distribution with one degree of freedom, according to Wilks’ theorem. The fitted models for comparative analyses were:

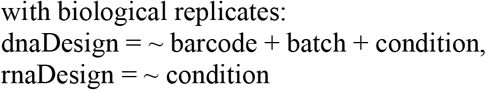

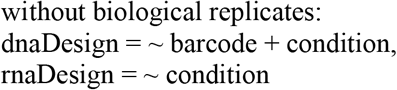

### Other statistical analyses

One-way analysis of variance and Spearman’s correlation tests were performed using Sidak’s adjustment for multiple comparisons^68^. Hypergeometric tests were performed using an online tool (https://systems.crump.ucla.edu/hypergeometric/).

Promoter rankings were given based on |log_2_ fold change| for promoters with an FDR < .05 in the all-replicate comparison and a minimum barcode number of 20. Heatmaps were generated using Morpheus^69^. Hierarchical clustering was performed via Morpheus using Euclidean distance with complete linkage. Dose response curves were generated using GraphPad Prism 9.5.0 using three or four parameter models.

## Data and code availability

Raw sequencing data will be deposited at NCBI prior to publication. Example datasets and our full analysis pipeline are available for immediate use at: https://github.com/JGEnglishLab/TRE-MPRA.

## Acknowledgements

We thank Brian Dalley, Opal Allen, and the University of Utah Huntsman Cancer Institute High-Throughput Genomics Core Facility for advice and use of their equipment for Illumina sample preparation and sequencing. This work was supported by the NIH NIGMS (DP2 GM146247).

