## Supplemental Figures 1-7 for "Discovery and Validation of Context-Dependent Synthetic Mammalian Promoters"

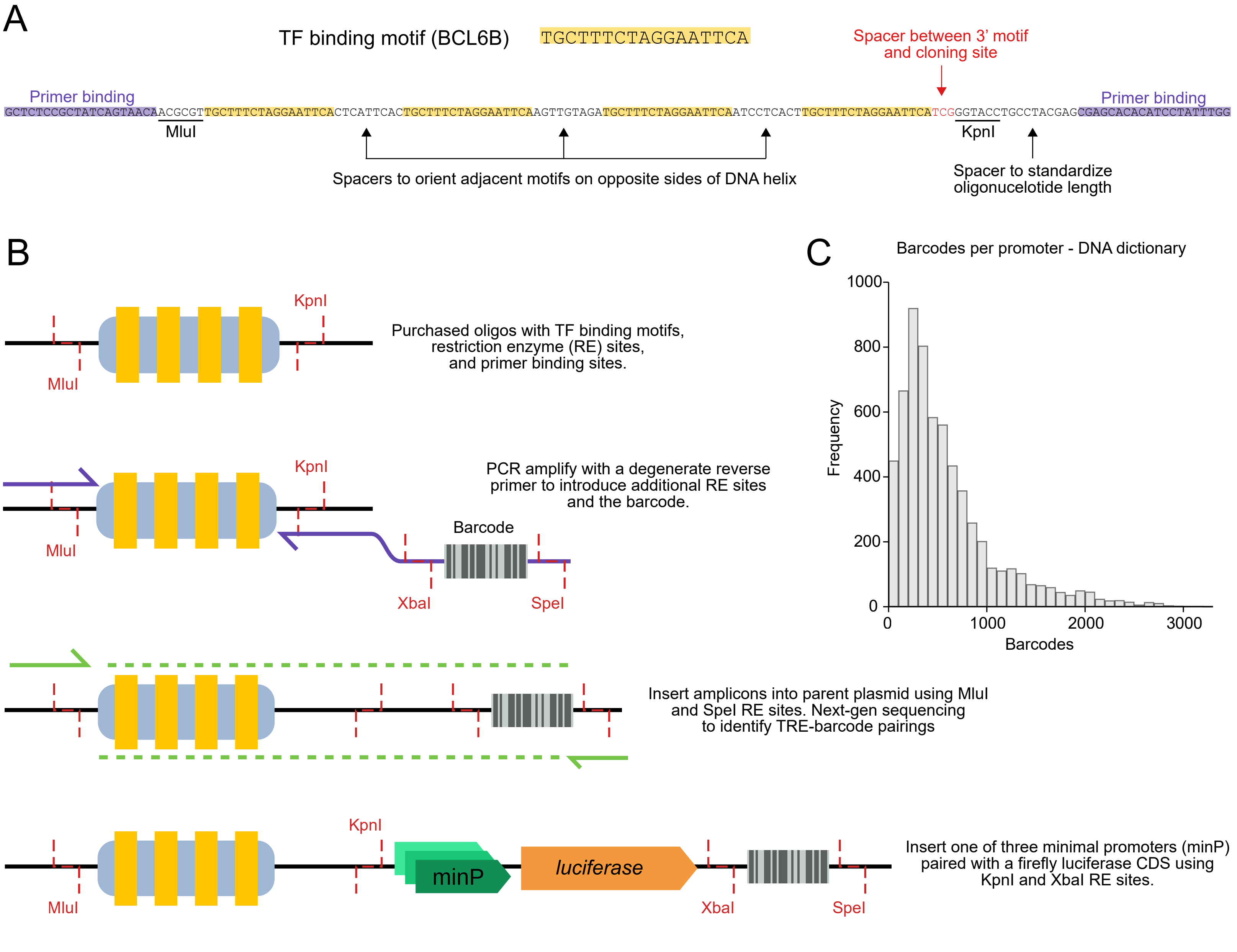


**Figure S1 - TRE unit design and promoter production. A**) Schematic of the TRE unit oligonucleotides used for MPRA library cloning. **B**) Graphical overview of TRE-MPRA library production. **C**) Histogram of the barcodes per promoter in the plasmid library barcode dictionary as determined by NGS.

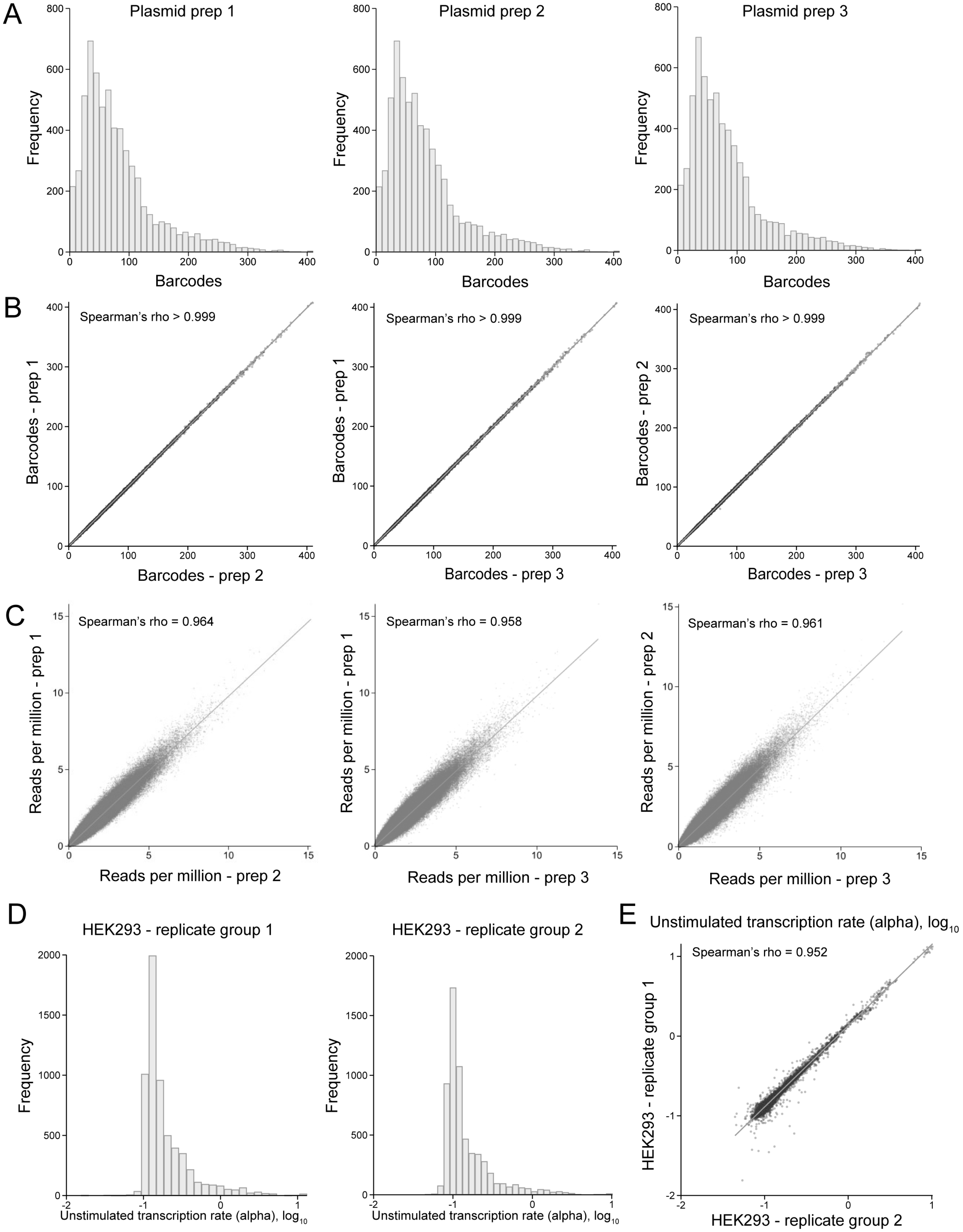


**Figure S2 - Comparison of independent TRE-MPRA replicates. A**) Histograms of the number of barcodes identified by NGS for each promoter. **B**) Pairwise comparisons of the number of barcodes detected for each promoter across plasmid preparations. **C**) Pairwise comparisons of the reads per millions for each barcode across plasmid preparations. **D**) Histograms of the estimated transcription rates (alphas) for promoters derived by MPRAnalyze in untreated HEK293 cell replicate groups. **E**) Scatterplot comparing promoter alphas between untreated HEK293 cell replicate groups.

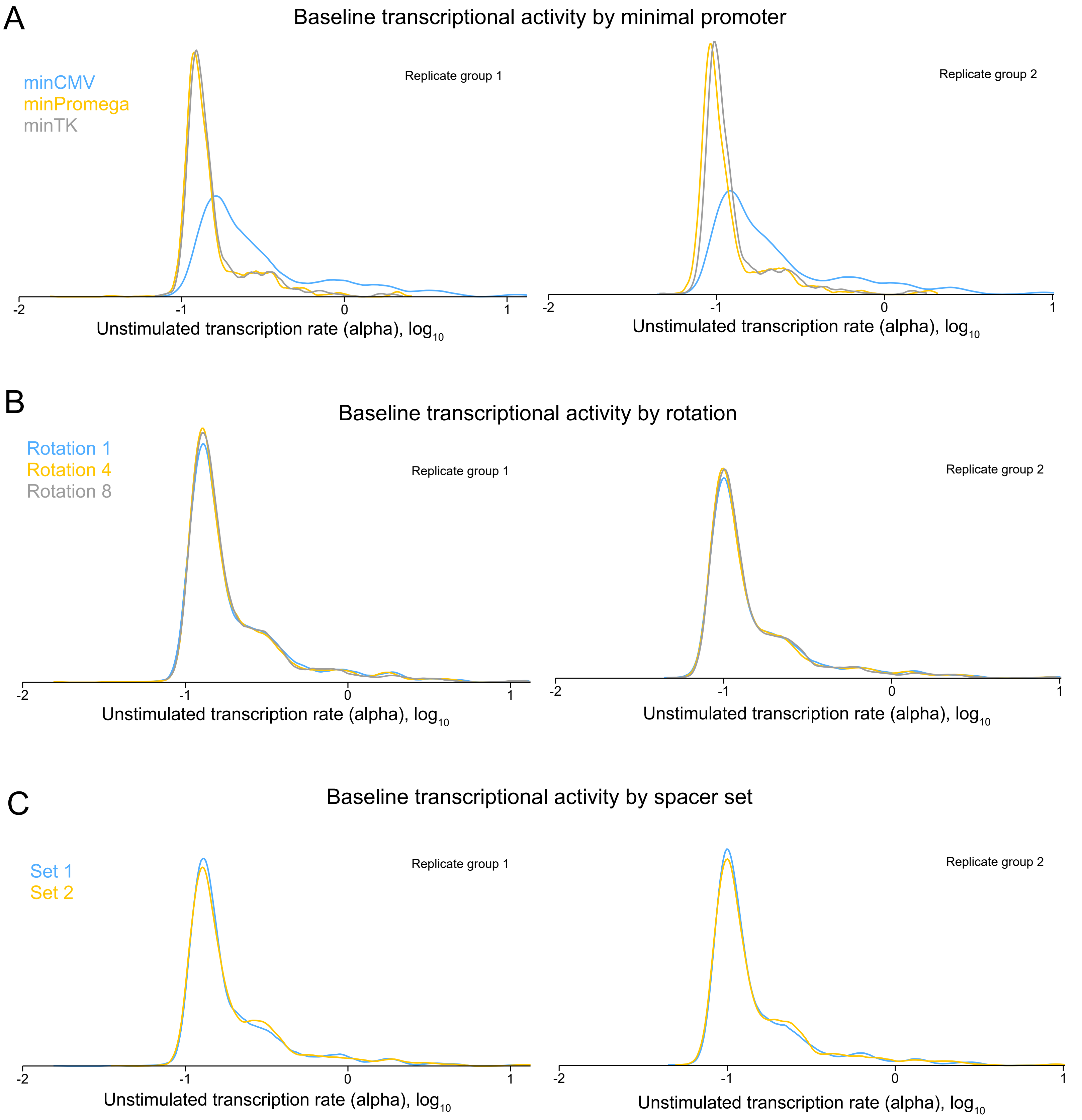


**Figure S3 - Effects of promoter design on transcription rates. A**) Density plots of baseline transcription rate estimates (alphas) for each minimal promoter across two independent sets of untreated HEK293 replicates. **B**) Density plots of baseline transcription rate estimates (alphas) for each nucleotide distance between TRE units and the minimal promoter (span) across two independent sets of untreated HEK293 replicates. Numbers shown (1,4,8) represent the nucleotide distance from the TRE unit and the minCMV or minTK promoters. Distances to the minProm are longer but maintain three rotations relative the DNA double helix. **C**) Density plots of baseline transcription rate estimates (alphas) for each of two spacer sets across two independent sets of untreated HEK293 replicates.

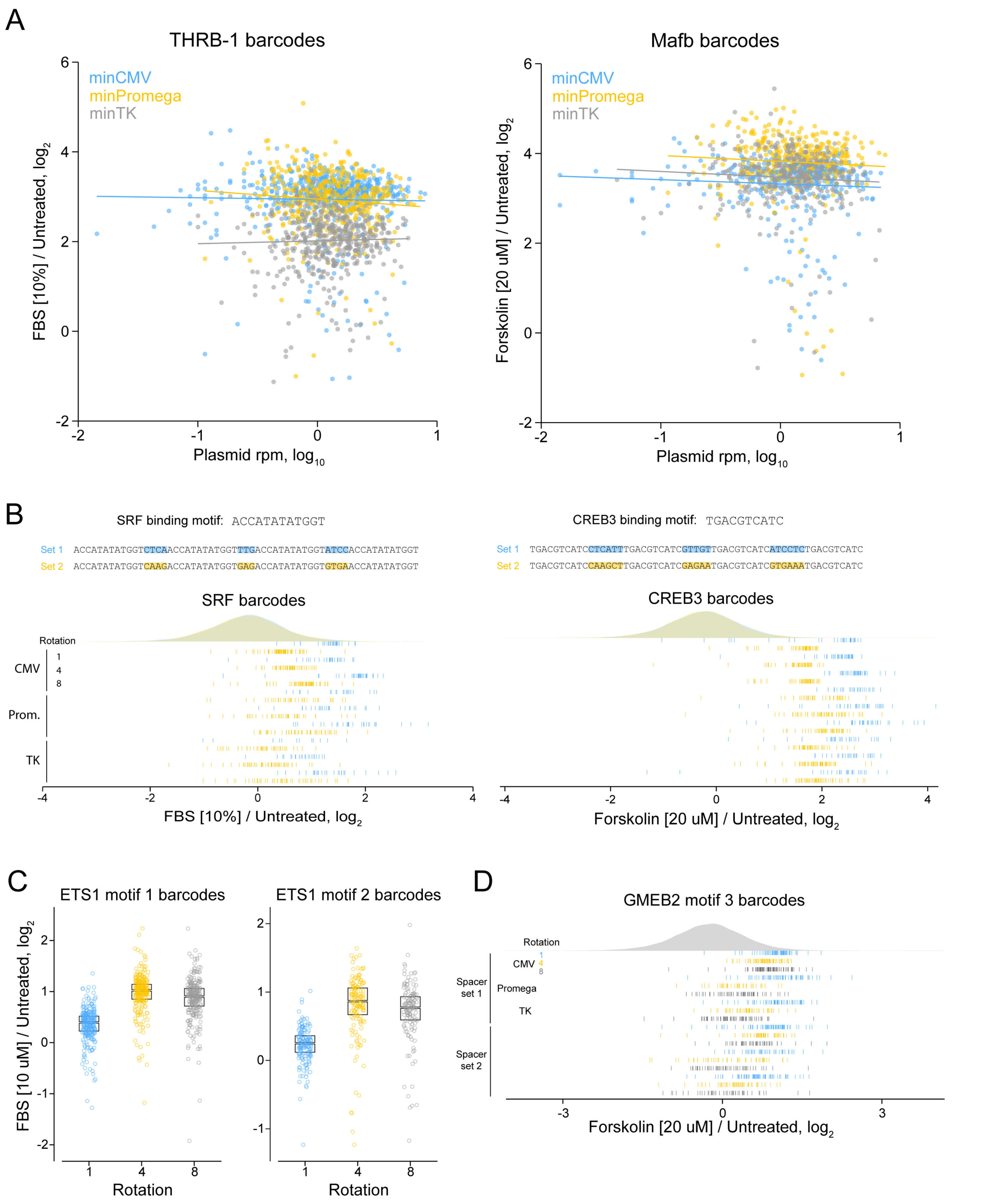


**Figure S4 - Promoter design effects on responses to FBS and forskolin. A**) Scatterplots comparing barcode representation in the plasmid library to the observed fold changes for THRB-1 (left) or Mafb (right) units. Colored lines indicate the linear fit of data in the displayed logs. There was no significant correlation between fold change and plasmid reads per million for any comparison. **B**) Raincloud plots for TRE units showing altered response to fetal bovine serum (left) or forskolin (right) depending on the spacer set incorporated. Each barcode is represented by a vertical line. The density plots indicate negative control barcodes with a given spacer set. **C**) Boxplots of barcode fold changes for two ETS1 motifs following FBS treatment according to the rotation relative to the minimal promoter. Boxes indicate the median and interquartile range. **D**) Raincloud plot of fold change following forskolin treatment for promoters containing the GMEB2 motif 3. Each barcode is represented by a vertical line. The density plot indicates negative control barcodes.

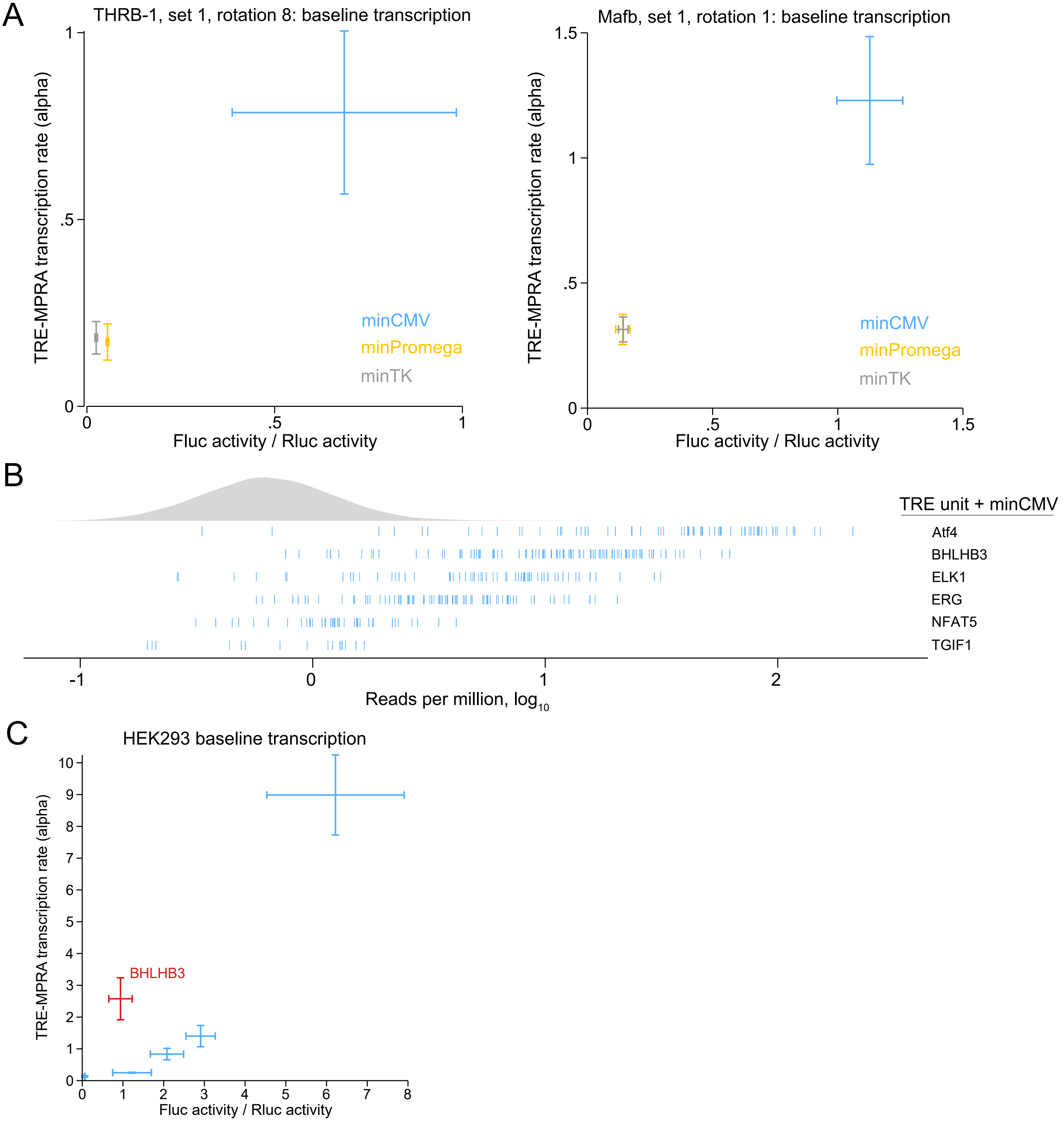


**Figure S5 - Orthogonal validation of baseline transcription activities in untreated HEK293 cells. A**) Comparisons of estimated transcription rates (alphas) by minimal promoter from the TRE-MPRA experiment and the observed raw luciferase ratios from orthogonal dual luciferase assays for the THRB-1 and Mafb units tested in Figure 2. Error bars indicate the standard deviation and their intersection indicates the mean. Alphas were derived from two independent experiments. Dual luciferase data are from three independent experimental replicates (n=4, N=3). **B**) Raincloud plot of the observed reads per million in untreated HEK293 cells for each barcode of six promoters. The gray density plot represents the barcodes of negative controls. **C**) Comparisons of estimated transcription rates (alphas) from the TRE-MPRA experiment and the observed raw luciferase ratios from orthogonal dual luciferase assays for six promoters shown in B. Error bars indicate the standard deviation and their intersection indicates the mean. Alphas were derived from two independent experiments. Dual luciferase data are from three independent experimental replicates.

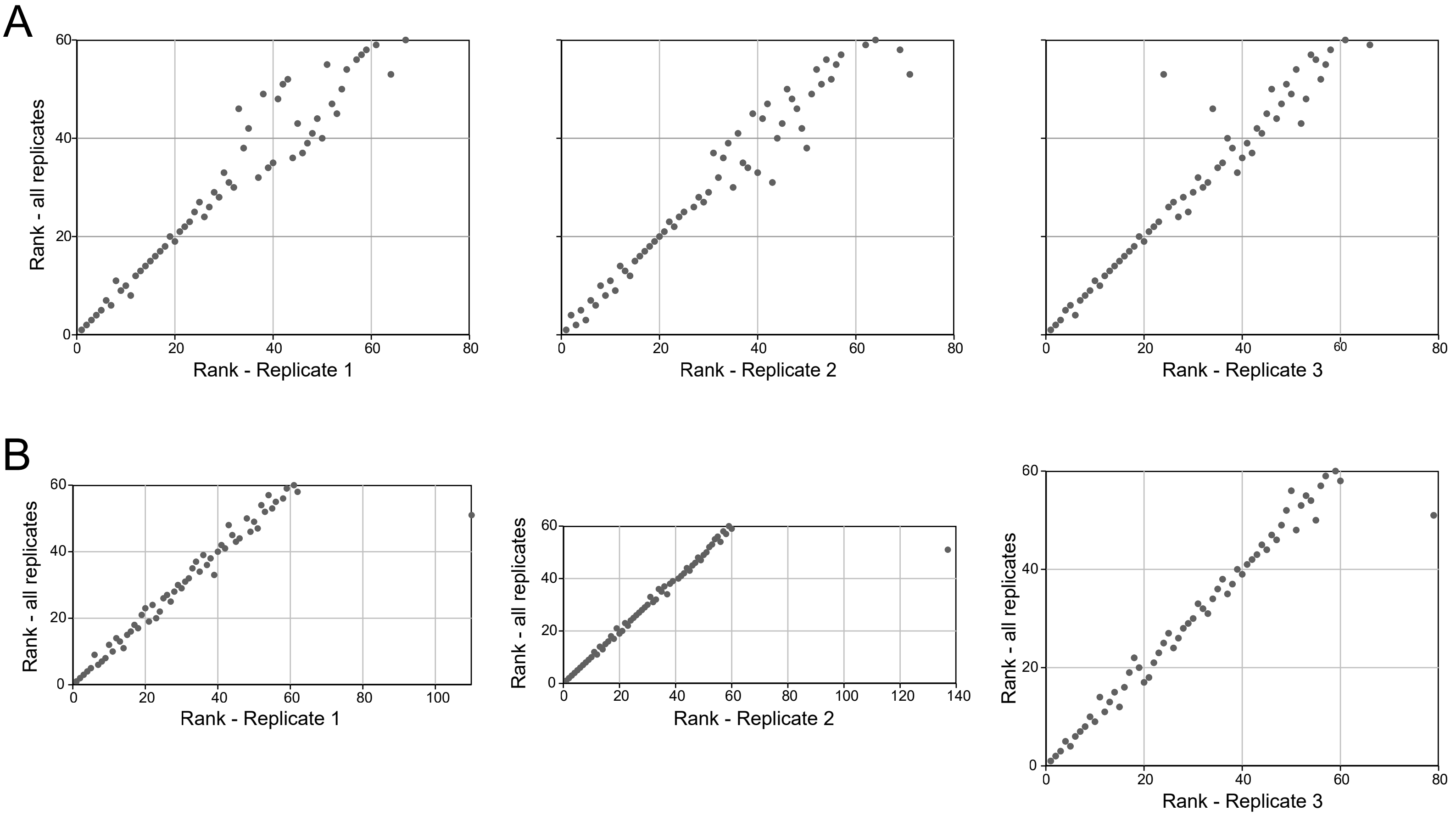


**Figure S6 - Comparing promoter rankings by single or multiple experimental replicates. A)** Scatterplots comparing the rankings of fold change absolute values of promoters following FBS treatment in HEK293 cells. Each individual replicate is compared to the complete group of three replicates. **B)** Scatterplots comparing the rankings of fold change absolute values of promoters following forskolin treatment in HEK293 cells. Each individual replicate is compared to the complete group of three replicates.

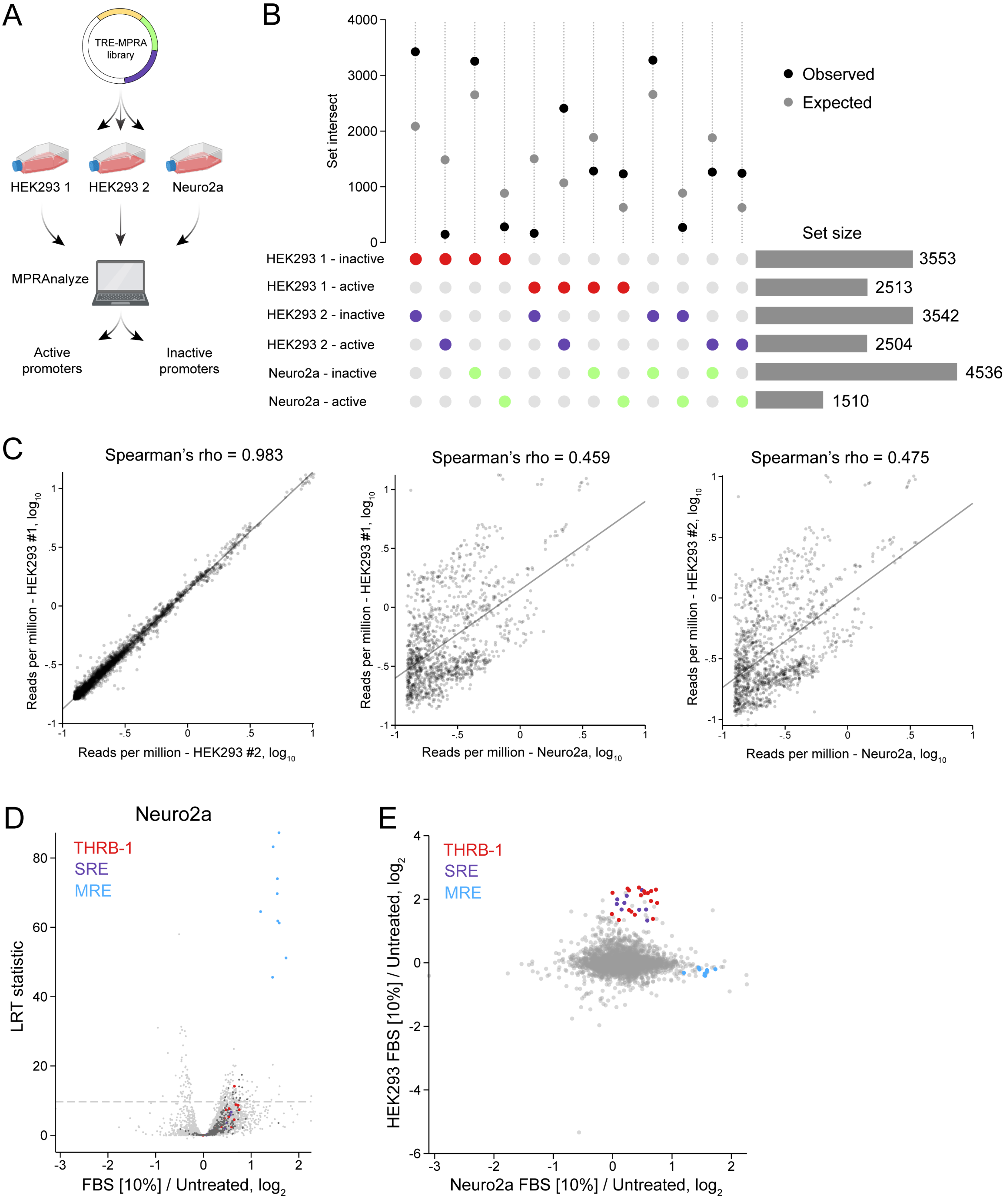


**Figure S7 - Cell background-specific TRE activities. A**) TRE activity classification using MPRAnalyze. **B**) Upset plot comparing the sets of active and inactive promoters from two independent HEK293 datasets and from Neuro2a cells. Sets being compared pairwise are indicated by colored circles. All set intersections were significantly different from what was expected by chance according to the hypergeometric test ($P < .001$). **C**) Scatterplots comparing the RNA reads per million for promoters classified as active between cell types. **D**) Volcano plot of promoter responses following stimulation of Neuro2a cells with 10% FBS for six hours, in comparison to untreated cells. The dashed line indicates an FDR threshold of 5%. Negative control promoters are indicated by dark gray data points. **E**) Scatterplot comparing the fold changes observed following FBS treatment in HEK293 and Neuro2a cells.
